# 3D imaging and morphometry of the heart capillary system in spontaneously hypertensive rats and normotensive controls

**DOI:** 10.1101/2020.08.05.237487

**Authors:** Camilla Olianti, Irene Costantini, Francesco Giardini, Erica Lazzeri, Claudia Crocini, Cecilia Ferrantini, Francesco Saverio Pavone, Paolo Guido Camici, Leonardo Sacconi

**Affiliations:** European Laboratory for Non-Linear Spectroscopy, Florence, Italy; National Institute of Optics, National Research Council, Florence, Italy; University of Colorado Boulder, Boulder, USA; University of Florence, Florence, Italy; Università Vita-Salute San Raffaele, Milan, Italy; Institute for Experimental Cardiovascular Medicine, University of Freiburg, Freiburg, Germany

**Author notes:** These authors contributed equally (Camilla Olianti, Irene Costantini).

## Abstract

Systemic arterial hypertension is a highly prevalent chronic disease associated with hypertensive cardiomyopathy. One important feature of this condition is remodelling of intramural small coronary arteries and arterioles. Here, we investigated the implications of this remodelling in the downstream vascular organization, in particular at the capillary level. We used Spontaneously Hypertensive Rats (SHR) exhibiting many features of the human hypertensive cardiomyopathy. We generated 3D high-resolution mesoscopic reconstructions of the entire network of SHR hearts combining gel-based fluorescent labelling of coronaries with a CLARITY-based tissue clearing protocol. We performed morphometric quantification of the capillary network over time to assess capillary diameter, linear density, and angular dispersion. In SHRs, we found significant remodelling of the capillary network density and dispersion. SHR capillary density is increased in both ventricles and at all ages, including before the onset of systemic hypertension. This result suggests that remodelling occurs independently from the onset of systemic hypertension and left ventricular hypertrophy. On the contrary, capillary angular dispersion increases with time in SHR. Consistently, our multi-modal imaging underlined a strong correlation between vascular dispersion and cellular disarray. Together our data show that 3D high-resolution reconstruction of the capillary network can unveil anatomic signatures in both physiological and pathological cardiac conditions, thus offering a reliable method for integrated quantitative analyses.

## INTRODUCTION

In the heart, large epicardial coronary conductive arteries (diameter >500 μm) branch into prearterioles (diameter 500–100 μm), intramyocardial arterioles (diameter <100μm) and finally, capillaries. Numerous bifurcations and anastomoses between capillaries create a dense vascular network intimately entwining cardiomyocytes allowing the diffusion of nutrients and oxygen. Microvascular remodelling is a process consisting of structural changes at the level of both systemic and coronary arterioles which has been described in patients with increased cardiac pressure overload consequent to sustained (untreated) systemic hypertension^1,2^. Arteriolar remodelling is characterized by vessel wall thickening as a result of hypertrophy of smooth muscle and increased collagen deposition ^3^.

However, whether microvascular remodelling precedes or follows the onset of systemic hypertension and left ventricle (LV) hypertrophy and how it correlates with other elements of myocardial remodelling (e.g. myocyte disarray) remains incompletely understood.

The Spontaneously Hypertensive Rat (SHR) model recapitulates many features of the human phenotype ^4^ ^5^. Patients with essential arterial hypertension demonstrate abnormal vasodilator capacity either during increased cardiac metabolic demand or during pharmacological vasodilation ^6^. Previous studies have shown that the medial area of arterioles is increased in SHR as compared to control counterparts already at 4 week of age, when arterial blood pressure was unchanged ^3^. This remodelling of the resistance vessels (the arterioles) increased progressively with aging in terms of medial area increase and decrease of lumen/vessel area.

Technical limitations characterize the bi-dimensional and non-integrated assays often used for studying microcirculation remodelling in hypertension. Bi-dimensional histo-morphometric analyses employing standard eosin-hematoxylin staining have provided both quantitative and qualitative information of capillary structure, but have shown poor reliability^7^. In fact, some reports that analysed vascular morphometries have indicated increased capillary density in hypertension compared to normotensive counterparts, suggesting compensatory angiogenesis in response to hypoxia, while some others have shown capillary rarefaction in SHR hearts, as a consequence of hypertension. To overcome bi-dimensional artefacts, several techniques for 3D anatomical assessment have been developed. Among them, Vascular Corrosion Casting (VCC) allows for 3D visualization of the microvasculature, including capillaries, using TEM microscopy ^8^. This technique enables high-resolution 3D reconstructions of the network; however, it confines the study to reduced volumes. Other approaches have exploited optical sectioning of fluorescent confocal microscopy to image brain and cardiac vascularization in rodents ^9^ ^10^, leading to 3D acquisitions of tissue sections with a micrometer resolution. Though, optical imaging is considerably limited by light scattering, a physical phenomenon that prevents light to penetrate deep into the tissue ^11^. The CLARITY protocol ^12^, in combination of 2,2’-Thiodiethanol (TDE) ^13^, a refractive index-matching agent, has proven to effectively clarify murine cardiac tissues ^14^, significantly reducing light scattering and preventing structural deformations of the specimen. However, 3D reconstruction of capillaries of cleared hearts is still ongoing. Recently, Di Giovanna et al. combined vascular perfusion with the BSA-FITC gel ^15^ with CLARITY/TDE clearing to perform whole mouse brain vasculature reconstruction, with remarkable signal-to-noise ratio and a high-quality imaging of the specimen.

In the present work, we overcame the restrictions imposed by current techniques for cardiac vasculature analysis, developing a protocol to successfully investigate capillary remodelling in SHR hearts through 3D structural reconstructions of large volumes with non-linear optical microscopy. Our work focused not only on studying of morphometric alterations of the capillary network in hypertensive conditions, but also on identifying the temporal correlation between the remodelling and the onset of arterial hypertension.

## RESULTS

### Vasculature staining in cleared rat heart

To investigate the microcirculation architecture, we developed a new approach to achieve 3D reconstruction of the coronary system and assess capillary remodelling. We optimized a staining protocol based on vascular perfusion with BSA-FITC gel. The gel is characterized by a low viscosity at temperature > 37°C and quickly solidifies at 4°C. The fluorophore (Fluorescein isothiocyanate, FITC) is conjugated to the high molecular-weight protein Bovine Serum Albumin (BSA) that prevents extravasation and confines the fluorescent staining to the vessels. Hearts were explanted and perfused with the gel at body temperature, then the gel solidification was favoured by keeping the heart in contact with ice-cold water. After complete solidification of the gel, hearts were optically cleared using a modified CLARITY protocol (*Figure 1A*). Finally, to achieve transparency, hearts were incubated in increasing concentration of 2,2’-Thiodiethanol: an agent with the same refractive index of biological samples, thus reducing light scattering during image acquisition (*Figure 1B*). This BSA-FITC/CLARITY protocol produces a homogeneous staining of capillaries enabling the 3D reconstruction of the microcirculation using Two-Photon Fluorescence Microscopy (TPFM) (*Figure 1C and Supplementary movie*). To verify that the protocol did not introduce systematic alterations to microcirculation morphometries, we analysed the capillary network in acute *ex-vivo* experiments where explanted hearts were stained with Dextran Fluorescein to label vessels, but were not clarified. We promptly imaged right and left ventricles, acquiring 50 μm deep stacks from the heart surface due to light scattering in non-clarified tissues. Vascular morphometries were analysed and compared to those of clarified samples (in this analysis, we included only the first 50 μm of the stacks). Capillary mean diameter, density, and angular dispersion were not different between *ex-vivo* and clarified hearts. (*Figure 1D*).

**Figure 1:**
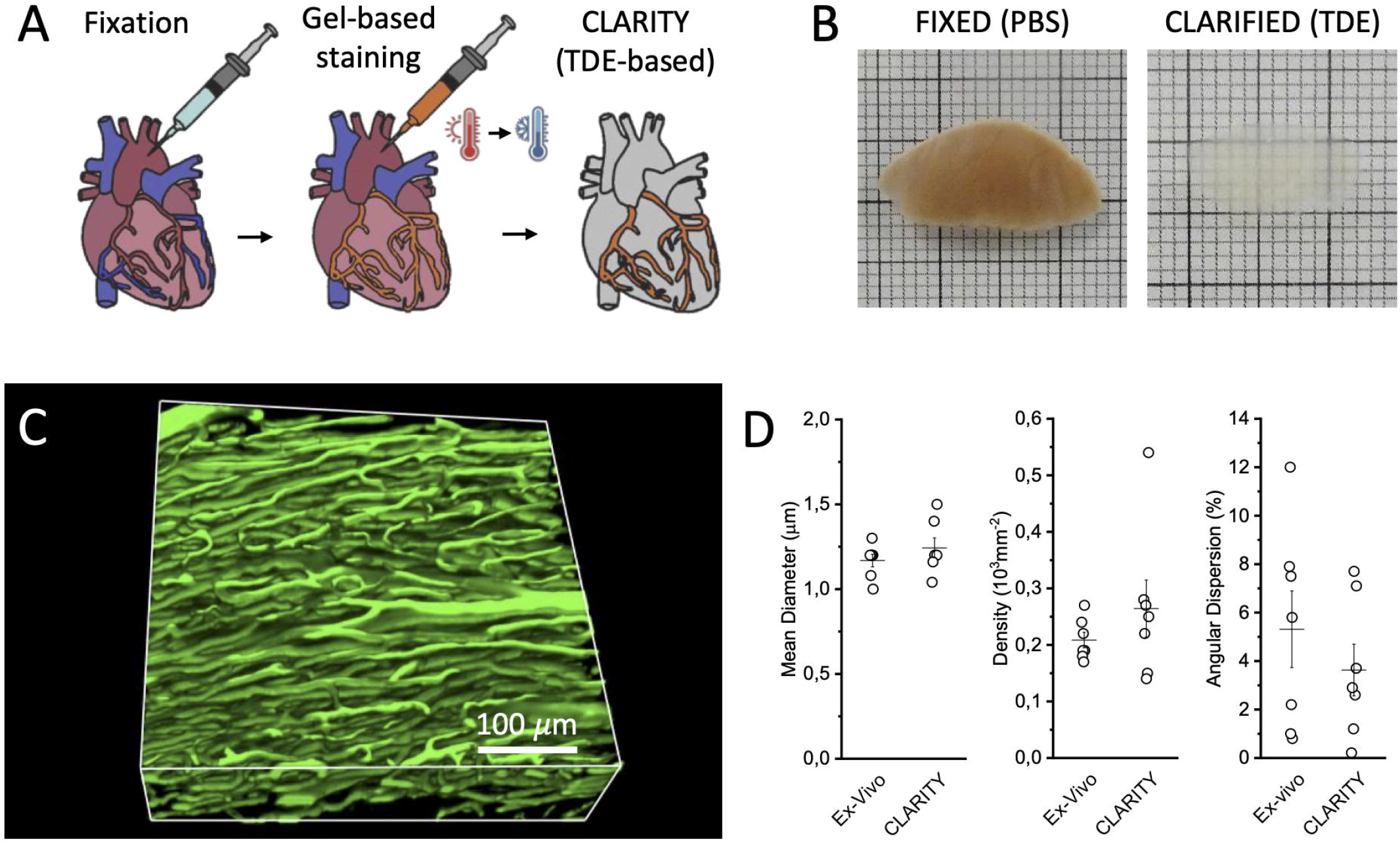
3D imaging of the coronary microcirculation of clarified hearts. A. Diagram showing the main steps of the BSA-FITC/CLARITY protocol. Heart is rapidly isolated, cannulated through the proximal aorta and perfused with fixative solution. Coronaries are stained with BSA-FITC gel and the organ is subjected to modified CLARITY protocol. B. Section of a right ventricle after fixative solution perfusion in PBS (left) and after TDE clearing (right). C. Representative 3D reconstruction of a stack of 450 × 450 × 300 µm^3^ acquired with a custom-made Two Photon Fluorescence Microscope. 3D reconstruction was created using ImageJ 2.0.0-rc-71/1.52p (https://fiji.sc). D. Validation of the sample preparation protocol: morphometric parameters obtained in acute experiment of ex-vivo hearts (Ex-Vivo) and in BSA-FITC cleared samples (CLARITY). Student T-test applied, p-value = 0.30; 0.31; 0.39. Plots were created using OriginPro 9.0 (https://www.originlab.com).

### Morphometry of coronary microcirculation

We employed the BSA-FITC/CLARITY protocol on right and left ventricles of SHR and normotensive Wistar Kyoto (WKY) animals at different ages: 4, 8, 18 and 24 weeks old in line with previous investigation on SHR microvascular remodelling ^3^. Capillary mean diameter, density, and angular dispersion were assessed from morphometries for each experimental group. The mean capillary diameter in both right and left ventricles of SHR compared to age-matched normotensive controls does not show any specific trend (*Figure 2B*). Capillary density is significantly increased in both RV and LV of SHR already at 4 weeks of age compared to WKY, when systolic blood pressure is still comparable between SHR and WKY (*Figure 2C*). Increased capillary density is maintained with aging in SHR as compared to WKY and does not change over time within the same experimental group, suggesting that those abnormalities are early manifestations of microcirculation remodelling in SHR hearts. Differently, capillary angular dispersion is increased in SHR with a clear time dependent rising effect (*Figure 2D*) suggesting a connection between capillary disorganization and tissue remodelling occurring in systemic hypertension ^16^. Increased vascular density and tortuosity can be appreciated in the Maximum Intensity Projection (MIP) of the stacks (*Figure 2A*).

**Figure 2:**
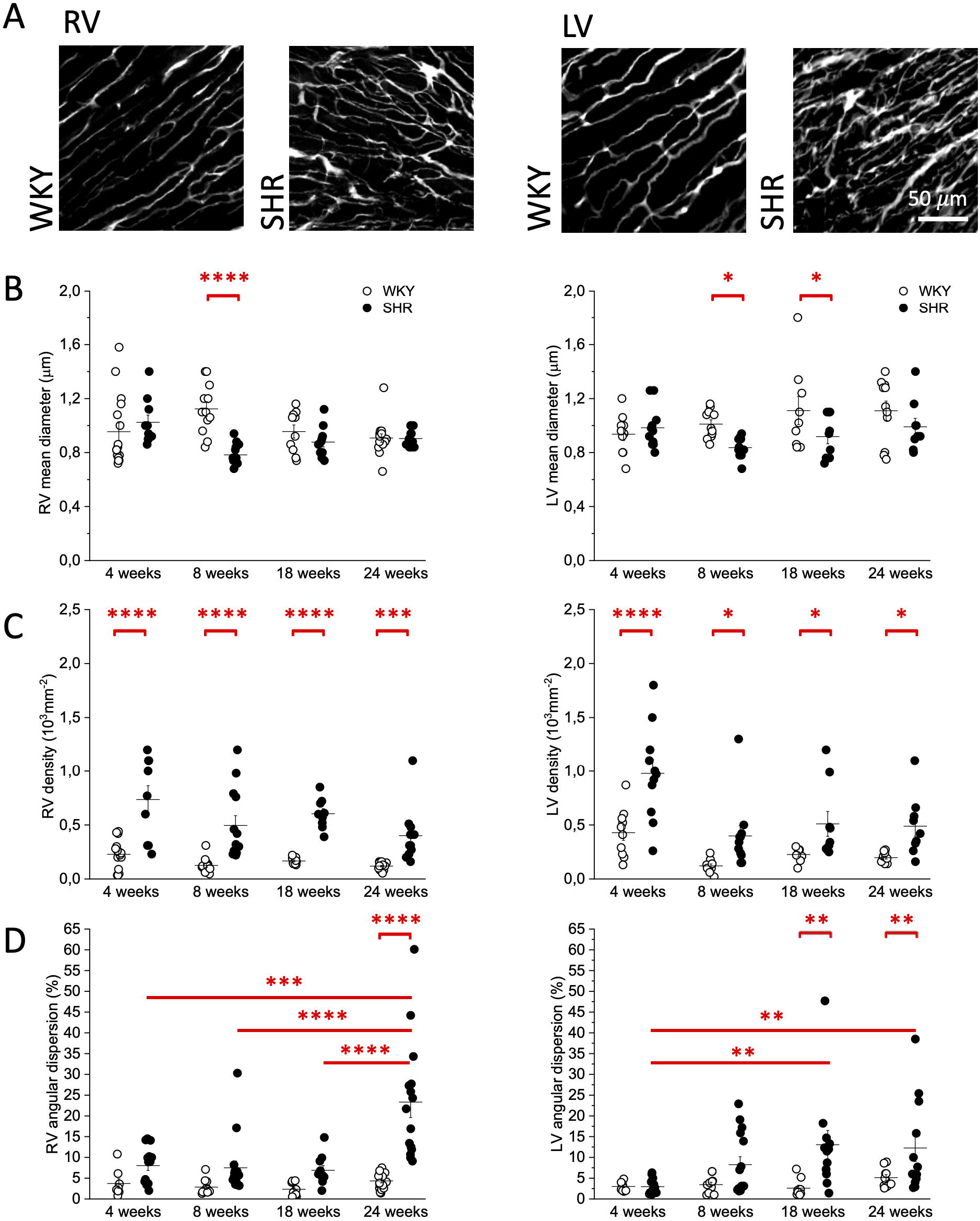
Morphometry of the coronary microcirculation in Spontaneously Hypertensive Rats and normotensive controls. Microcirculation morphometries of Right (RV) and Left (LV) ventricles: A. Representative maximum intensity projection of 60 µm depth of the microcirculation in WKY and SHR. Maximum intensity projections were created using ImageJ 2.0.0-rc-71/1.52p (https://fiji.sc). B., C., and D. Mean lumen diameter, capillary density, and angular dispersion of the capillary network in RV and LV of WKY and SHR at different time points of age. Two-way ANOVA with fisher LSD Post Hoc test: p<0.05 *; p<0.005 **; p<0.001 ***; p<0.0001 ****. Plots were created using OriginPro 9.0 (https://www.originlab.com).

### Multi-modal imaging for tissue cytoarchitecture characterization

To correlate microcirculation abnormalities to tissue cytoarchitecture changes, we stained samples, previously treated with the BSA-FITC/CLARITY protocol, with Wheat Germ Agglutinin (WGA) conjugated with Alexa Fluor 594 to mark cellular membrane. We then investigated cellular arrangement by two-photon fluorescence imaging and collagen deposition through the second-harmonic generation (SHG) signal. A representative image of the multi-modal analysis is shown in *Figure 3A*.

**Figure 3:**
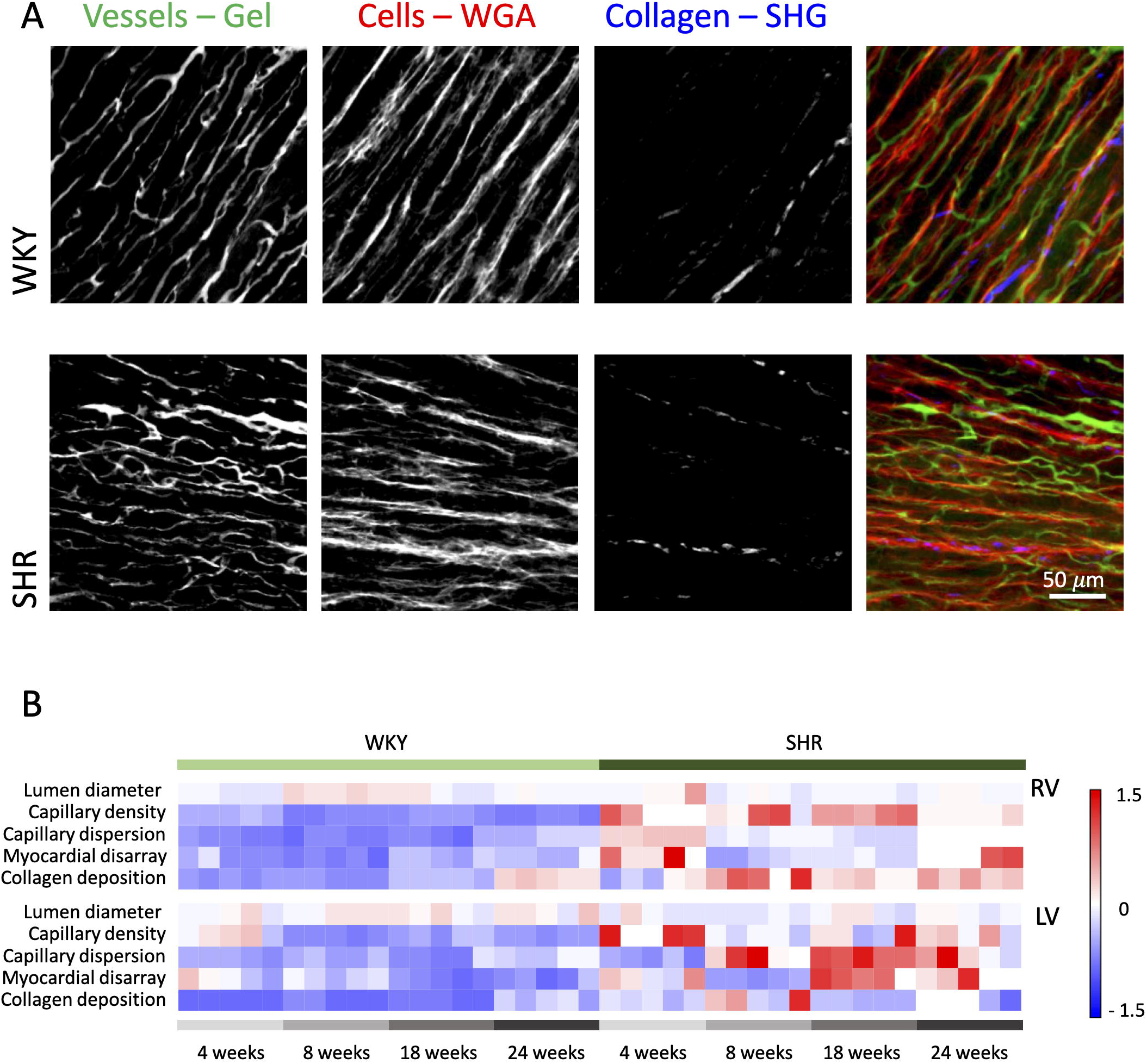
3D multi-modal imaging and quantification. A. Representative maximum intensity projection of 60 µm deep stack of WKY and SHR ventricles showing from the left: capillaries labelled with BSA-FITC gel, negative image of cells traced with WGA-Alexa Fluor 594, collagen deposition imaged by second-harmonic generation, and merge of the three different channels (vessels in green, cells in red, collagen in blue). Maximum intensity projections were created using ImageJ 2.0.0-rc-71/1.52p (https://fiji.sc). B. Heat Map summarizing vascular, cellular and collagen parameters in all the experimental classes. Five different ventricular regions are shown for each experimental class. All the parameters have been normalized on the same scale. RV is right ventricle (above) and LV is left ventricle (below). Heat Maps were created using OriginPro 9.0 (https://www.originlab.com).

Vascular, cellular and collagen parameters of the different experimental classes are summarized in the Heat Map of *Figure 3B*. Myocardial disarray is significantly different in SHR compared to WKY predominantly in LV and at 18 and 24 weeks, while collagen deposition is increased in SHR compared to controls in all the time-points.

To study the relationship between vascular remodelling and tissue cytoarchitecture we performed a Pearson’s correlation analysis between capillary density or angular dispersion with myocardial disarray or collagen content (*Figure 4*). Cellular disarray shows a significant correlation with capillary angular dispersion in line with the hypothesis that capillary misalignment is interconnected with tissue remodelling. No correlation is found between capillary density and myocardial disarray in agreement with the early manifestation of the capillary density growth found in *Figure 2C*. Finally, collagen content shows no correlation with vascular morphometries, thus suggesting that capillary abnormalities are independent from collagen features in SHR hearts.

**Figure 4:**
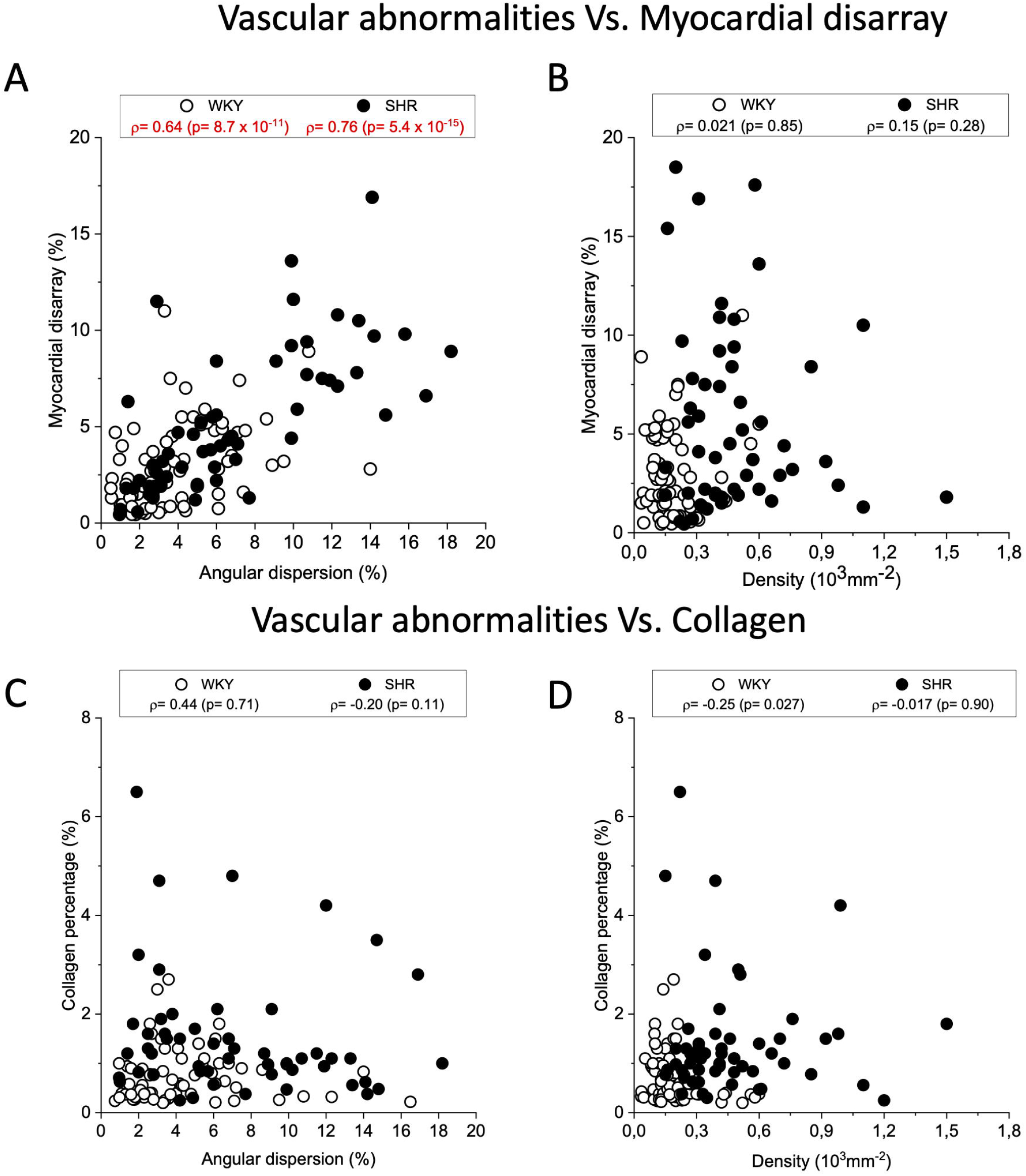
Vascular disorganization correlate with myocardial disarray. Correlation analysis between vascular morphometries and tissue cytoarchitecture in WKY and SHR ventricles. A. angular dispersion and cellular disarray, B. vasculature density and cellular disarray, C. angular dispersion and collagen percentage, and D vasculature density and collagen content (%). Plots were created using OriginPro 9.0 (https://www.originlab.com).

## DISCUSSION

To date, the temporal relationship between vascular remodelling and the onset of hypertrophy remains controversial. Capillary proliferation has been widely considered as an adaptive response to myocardial hypertrophy, however novel evidences show that an increase in capillary mass *per se* can promote hypertrophic growth of myocytes ^2^. In this work, we imaged and quantified the 3D cardiac capillary structure and we described its remodelling in the Spontaneously Hypertensive Rats (SHR) over-time. To this aim, we generated 3D capillary morphometries of SHR RV and LV, and analysed mean lumen diameter, density and angular dispersion over-time, from 4 to 24 weeks old animals. Our 3D analyses highlighted no clear trend in lumen diameter between SHR and WKY, while a significant increment of capillary density in SHR hearts as compared to age-matched WKY controls was observed even before the onset of the disease, i.e. at 4 weeks of age. In particular, capillary density in SHR hearts is at least two-fold higher than WKY controls. This difference did not depend on age within the experimental groups. Conversely, age-dependent effects were observed on capillary angular dispersion, especially in the LV. To better understand the anatomical context in which the observed remodelling takes place, we performed multi-modal imaging to analyse myocardial disarray and collagen deposition. We found a significant increase of myocardial disarray at later time-points (18-24 weeks) in LV similarly to capillary angular dispersion, suggesting a connection between capillary disorganization and tissue remodelling in systemic hypertension over-time. A Pearson’s correlation analysis between vascular remodelling and myocardial disarray confirmed the presence of a strong correlation.

Our findings suggest that the increased capillary density observed at early stage in SHR hearts was an early feature of hypertensive cardiomyopathy that preceded the onset of hypertension (8 weeks old) and LV hypertrophy (12 weeks old) ^16^. These findings, in combination with the lack of differences between right and left ventricle capillary density, prompts us to speculate that capillary proliferation did not represent a hemodynamic response to increased systemic blood pressure or impaired arteriolar vasomotion, but rather was driven by genetic and local factors. On the contrary, we demonstrated that capillary disarray was a time-dependent process and, more importantly, it correlated with myocardial disarray, suggesting a vascular involvement of tissue reorganization in cardiac hypertrophy. These results could not be achieved with conventional bi-dimensional analyses. Three-dimensionality is essential to perform reliable quantitative analyses of the capillary network, whose morphology could be affected by the sectioning plane of the sample ^7^. To this aim, we developed a new successful approach to perform macroscopic reconstructions with a sub-cellular resolution using a clearing-based technology. The BSA-FITC/CLARITY methodology combined gel-based fluorescent staining of vasculature with a TDE-based optical clearing protocol. This approach offered great advantages in comparison to vascular endothelium immuno-staining since the gel fills the entire lumen, providing a great signal-to-noise ratio ^15^. The TDE-based CLARITY methodology was optimized for rat hearts, achieving full transparency of the tissues, thus reducing light scattering and improving image acquisition with two-photon fluorescence microscope. In addition, our approach allowed us to simultaneously investigate different components of the tissue permitting a multi-modal analysis of the specimen.

The 3D reconstruction of large volumes leads to the generation of an extensive amount of data that cannot be manually analysed by the operator, requiring a proper automatic analysis of 3D images ^17^. We optimized a pipeline to extract morphometric information as lumen diameter and density from segmented images; furthermore, we obtained vessels and cellular orientation by means of a software able to trace biological feature directionality in 3D through the generation of a map of versors.

The developed methodology can be used to investigate the role of the coronary architecture in the onset and progression of different pathologies besides hypertension. With little optimization, the protocol could be applied to other organs or species. Moreover, the combination of this new approach with Light-Sheet Fluorescence Microscopy will allow volumetric reconstruction of the whole coronary system of intact organs. Finally, the structural analyses obtained with our pipeline could be coupled to functional studies to unveil the relation that binds the anatomical organization of the coronary system to the cardiac muscle functions. We believe that the approach presented in this work will pave the way to new frontiers for anatomical studies and will allow to obtain a more complete understanding of the effects of therapeutics on the 3D structural organization of the tissues.

## METHODS

### Animal models

Male Wistar-Kyoto (WKY) and Spontaneously Hypertensive Rats (SHR) were analysed at different time-points of age: 4, 8, 18, and 24 weeks old. All animal procedures performed conform to the guidelines from Directive 2010/63/EU of the European Parliament on the protection of animals used for scientific purposes; experimental protocol is approved by the Italian Ministry of Health on July 6, 2015; authorization n° 944/2018-PR. All the animals were provided by Charles River, Milan, Italy.

### Heart isolation and perfusion

Animals were sacrificed while still beating under Isoflurane anaesthesia (3% Isoflurane in 100% oxygen at a flow rate of 1.0 L/min). The thorax was opened and heart was rapidly isolated. Heart has been cannulated through proximal aorta and perfused with 30 mL of Tyrode Solution (*Glucose 10mM, HEPES 10mM, NaCl 113 mM, MgCl_2_ 1,2 mM, KCl 4,7 mM; pH 7.4*) with a constant pressure of 10 mL/minute to remove blood from the vessels. Subsequently, the heart has been perfused with 24 mL of a fixative solution containing 4% of Paraformaldehyde (PFA) in Phosphate Buffered Saline (PBS) (pH 7.6). To perform Ex-Vivo experiments hearts were not perfused with the fixative solution and the vascular network has been immediately labelled after Tyrode Solution perfusion.

### Vascular network staining

Hearts were perfused with 30 mL of a fluorescent gel, containing FITC fluorophore (Fluorescein Isothiocyanate) conjugated with Bovine Serum Albumine (BSA), recently developed by Tsai et al ^18^ and previously used to stain brain capillary network ^15^. The gel contains 2% of Gelatine from Porcine Skin (no. G1890; Sigma-Aldrich) in PBS. Once melted, the solution was filtered, added with 0,05% of BSA-conjugated FITC (Albumin-fluorescein isothiocyanate conjugate, Sigma-Aldrich) and filtered again. The preparation was kept at 37°C in gentle shaking until use. Gel solidification into the vessels was favoured by keeping the organ in contact with iced water for 30 minutes.

For vascular network labelling of non-fixed hearts, the organ has been perfused with 50 mg of a dextran (Dextran fluorescein; Molecular Probes), solubilized in 2 mL of Tyrode Solution. Images were immediately acquired on left and right ventricles.

### Tissue clearing

Hearts were subjected to a tissue transformation protocol based on CLARITY methodology 12, modified to be used on cardiac tissue. Briefly, after vascular perfusion with BSA-FITC gel, tissues were incubated in 30 mL of PFA 4% in PBS O/N and the following day they were washed in PBS and sectioned in left atrium, right atrium, inter-ventricular septum, left ventricle and right ventricle. Left and right ventricles were subjected to CLARITY protocol: they were incubated in 30 mL of Hydrogel Solution *(4% Acryilamide, 0,05% Bis-Acryilamide, 0,25% Initiatior AV-044 in 0,01 M of PBS)* for 3 days at 4°C in gentle shaking. After 3 days the samples were de-gassed using a drier (KNF Neuberger, N86KT.18) and oxygen has been replaced by nitrogen. To favour gel polymerization, the samples have been kept at 37°C for 3 hours and then incubated in 30 mL of Clearing Solution (*Boric Acid 200 mM, 4% Sodium Dodecyl-Sulfate; pH 8,6*) at 37°C in shaking up to complete clearing of the tissue (about 12-16 weeks).

### Cellular membrane staining

After the clearing, the tissue has been washed in PBS at RT for 24 hours, then permeabilized with a PBS supplemented with 0.1% of Triton X-100 (Sigma-Aldrich) for 24 hours. The following day the sample has been stained with Wheat Germ Agglutinin (WGA) conjugated with Alexa Fluor 594 (W11262; dilution 1:100 in PBS-T; Invitrogen) for 24 hours.

### Refractive Index Matching with 2,2’-Thiodiethanol (TDE)

Samples were then cleared in increasing concentration of 2,2’-Thiodiethanol (TDE) in 0,01M PBS. The first two incubations at 20% and 47% were performed for 2 hours at room temperature, while the final concentration of 68% (TDE/PBS) was reached by incubating the sample for 12 hours at 37°C while gentle shaking.

### Image acquisition

For image acquisition we used a custom-made two-photon fluorescence microscope (TPFM) equipped with a mode locked Chameleon titanium sapphire laser (120 fs pulse width, 90 MHz repetition rate; Coherent Inc.); the laser operates at 780 nm and it has been coupled with a custom-made scanning system supported with a pair of galvanometric mirrors (LSKGG4/M; Thorlabs, Newton, NJ, USA). To acquire images the laser has been focused on the specimen with a refractive index tunable 25 × objective (LD LCI Plan-Apochromat 25x/0.8 Imm Corr DIC M27; Carl Zeiss, Oberkochen, Germany). A closed-loop xy stage (U-780 PILine XY Stage System; Physik Instrumente, Karlsruhe, Germany) allows radial displacement of the specimen and a closed-loop piezoelectric stage (ND72Z2LAQ PIFOC objective scanning system, 2 mm travel range; Physik Instrumente) allows shifting the objective along the Z-axis. Three independent GaAsP photomultiplier modules (H7422; Hamamatsu Photonics, Bridgewater Township, NJ, USA) acquire the fluorescent signal. Band-pass emission filters centred at 530 ± 55 nm and 618 ± 50 nm were used, respectively, for FITC and Alexa Fluor 594 detection, and a filter centred at 390 ± 18 was used for second-harmonic generation. Stacks of 450 × 450 μm, with a depth of 300-400 μm and a Z-step of 2 μm were acquired. For mosaic reconstruction serial Z-stacks of adjacent regions, laterally overlapped by 40 μm, were collected and images of corresponding Z-planes were stitched using the 3D stitching tool ZetaStitcher (https://github.com/lens-biophotonics/ZetaStitcher).

### Image analyses

Signals collected from the three different channels were split and images derived from each channel have been separately analysed.

Image resolution has been made isotropic: for such a purpose, ratio between the pixel-size in x-y (0,439 μm) and the one in z (2 μm) has been calculated, and the number of frames in Z has been rescaled through the resulting factor (4.55). For each stack, Mean Lumen Diameter and capillary Density have been automatically calculated using the software Amira 5.3 and a custom-made LabView interface (National Instruments). In details, for vascular network identification, we have applied Otsu algorithm on Amira, which automatically estimates a global threshold of the signal emitted by the vessels and then we performed a skeletonization of the network using the Autoskeleton algorithm. The output of the analysis does not consist in a single map of the system, but it is constituted by the sum of different segments. For each of them the software provides length, volume and mean radius. To estimate the vascular mean diameter of each stack, values obtained by the automatic segmentation of the vascular network have been divided into different groups, basing on the segments length (0 to 5 μm, 5 to 10 μm, 10 to 15 μm, 15 to 20 μm, 20 to 25 μm and 25 to 30 μm). Through the LabView Interface the Mode has been calculated for each group, finally, the Mean Diameter of the stack was calculated as the Mean value of the Modes. Density has been calculated as the ratio between the total vasculature surface and the total volume of the analysed stack. To validate the reliability of the Amira 5.3 software in calculating the morphometric parameters we performed a comparison between the mean radius value automatically calculated by the software Amira and the value obtained with a manual estimation (using the ImageJ software). We detected 25 different segments of 5 different stacks and we compared the value gained by Amira’s segmentation to that obtained manually tracing diameter in 10 different spots of each segment. The results verify the reliability of the automatic segmentation tool, the relationship between the two technique is linear as demonstrated by plotting the values on a graph and fitting them through the y = × line (*Supplementary Figure 1*). To quantify myocardial disarray and vascular angular dispersion in the tissue, we developed an automatic software tool to estimate the 3D orientation of vessels and cardiomyocytes. The software virtually dissects the volume in portions of about 60 μm. There, it applies a Structure Tensor (ST) analysis to extract the main direction of the cells or vessels, storing all these orientations as versors across the entire volume. Then, it estimates multiple local disarrays dividing the virtual architecture in portions of 120 μm. Disarray is defined as (1 – Alignment), where Alignment is the module of the average vector of local orientation versors. Finally, the global disarray of the entire volume is obtained averaging all local disarrays (*Supplementary Figure 2*).

To estimate collagen amount on total volume analysed, images have been rendered binary using the default ImageJ algorithm and the percentage of signal pixels on the total pixel amount has been calculated by dividing the amount of signal pixel for the total amount of pixels in the volume.

### Data analysis

Graphs and data statistical analyses were achieved using OriginPro 9.0 (OriginLab Corporation). We performed two-way ANOVA analysis to assess significant variance between both the two different experimental groups (WKY and SHR) and the four different time-points of age (4, 8, 18 and 24 weeks); Fisher LSD has been used to perform Post Hoc Analysis. For the correlation analyses between myocardial disarray, collagen percentage and vascular morphometries, we performed a Pearson’s Correlation test.

## FUNDING

This work was supported by the European Union Horizon 2020 research and innovation program under grant agreement [654148: Laserlab-Europe]; by the Italian Ministry for Education, University and Research in the framework of the Flagship Project [NANOMAX]; by Ente Cassa Risparmio di Firenze [WHOLE-HEART]; and by FAS-Salute [ToRSADE project]. C.C was supported by Human Frontiers Science Program fellowship (LT001449/2017-L) and by the American Heart Association (AHA) postdoctoral fellowship (20POST35211113).

## CONFLICT OF INTEREST

The authors declare no competing interests.

## DATA AVAILABILITY

All data generated or analysed during this study are included in this published article (and its Supplementary Information files).

## Notes

### Competing Interest Statement

The authors have declared no competing interest.

