## Supplementary Information for "3D imaging and morphometry of the heart capillary system in spontaneously hypertensive rats and normotensive controls"

*These authors contributed equally.

### SUPPLEMENTARY FIGURES

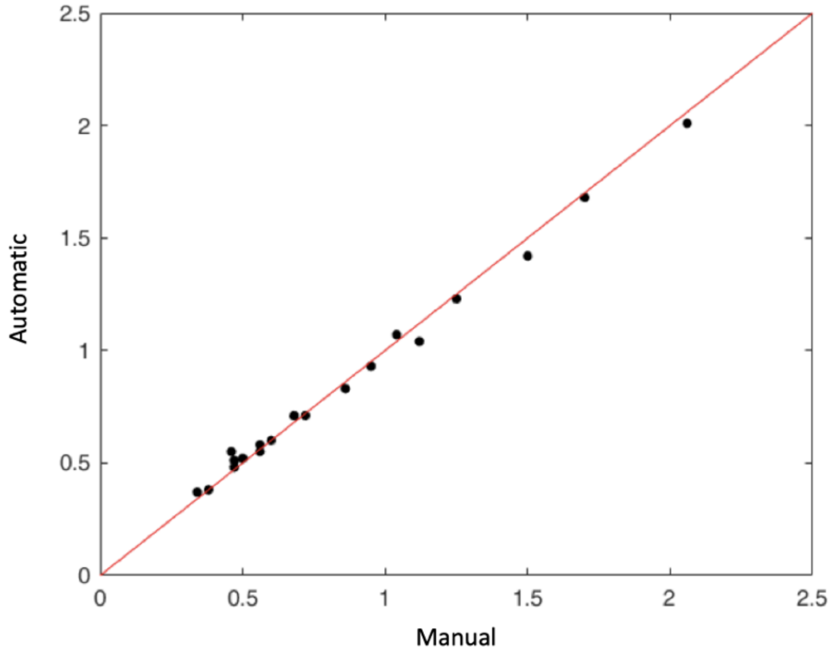


**Figure S1: Manual vs automatic estimation of mean lumen diameter**. The mean lumen diameter from 20 different segments has been estimated both manually (using ImageJ) and automatically (using Amira 5.3). The linear relationship between the manual and the automatic valuation of the mean lumen diameter validates the reliability of software Amira. Plot was created using OriginPro 9.0 (https://www.originlab.com).

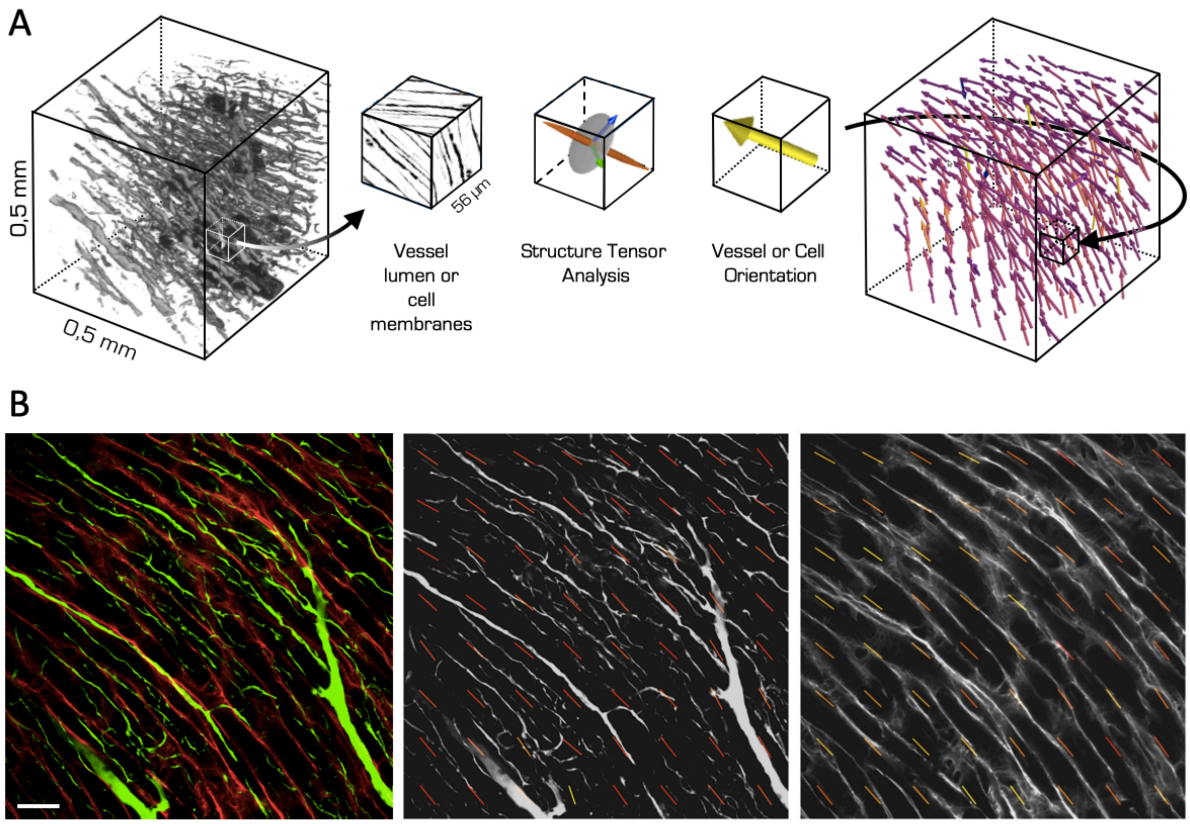


**Figure S2: Analysis of 3D local orientations of both vascular network and cardiomyocytes in rat heart tissues.** A. Scheme of the orientation analysis pipeline. The acquisition volume channels (red for cell membranes and green for vessels lumen) are analysed separately. For both channels, the volume is virtually dissected in portions of about 56um. Inside each portion, a Structure Tensor Analysis is applied directly on the grey levels. We collect the eigenvector with the smallest eigenvalue to estimate the mean direction of the signal gradient, i.e. the 3D orientation of local biological structures. Then, each orientation vector is re-allocated in a virtual volume to reconstruct the entire structural organization. The scheme of the orientation analysis pipeline was created with Microsoft Power Point 16.16.22 (https://www.office.com) using images created with ImageJ 2.0.0-rc-71/1.52p (https://fiji.sc). B. Example of the orientation analysis results. On the left, a representative (450 x 450) µm sample of right ventricle of a 24 weeks old WKY rat. In red, a single frame of WGA signal, with depth of 28 µm. In green, a Maximum Intensity Projection (MIP) of the first 56 µm in depth of the vessel stack. The 3D orientation vectors obtained analysing the vessel signal (in the centre) and the cell membranes (on the right) are superimposed on the original frame as 2D vectors (the colour represents the z components). Scale bar: 50 µm. MIPs were created using Python 3.6.8 (https://www.python.org).

### SUPPLEMENTARY MOVIE LEGEND

3D rendering of capillary network of a clarified WKY heart with vessels stained the BSA-FITC gel. 3D images were acquired using two-photon florescence microscope. Movie was created using Amira 5.3 (https://www.thermofisher.com/fr/en/home/industrial/electron-microscopy/electron-microscopy-instruments-workflow-solutions/3d-visualization-analysis-software/amira-life-sciences-biomedical.html).
